# Visualizing adenosine to inosine RNA editing in single mammalian cells

**DOI:** 10.1101/088146

**Authors:** Ian A. Mellis, Rohit K. Gupte, Arjun Raj, Sara H. Rouhanifard

**Affiliations:** Department of Bioengineering, University of Pennsylvania, Philadelphia PA; Genomics and Computational Biology Group, Perelman School of Medicine, University of Pennsylvania, Philadelphia PA

## Abstract

Conversion of adenosine bases to inosine in RNA is a frequent type of RNA editing, but important details about its biology, including subcellular localization, remain unknown due to a lack of imaging tools. We developed an RNA FISH strategy we called inoFISH that enables us to directly visualize and quantify adenosine-to-inosine edited transcripts *in situ*. Applying this tool to three edited transcripts (GRIA2, EIF2AK2 and NUP43), we found that editing of these transcripts is not correlated with nuclear localization nor paraspeckle association, and that NUP43 exhibits constant editing rates between single cells while the rates for GRIA2 vary.

---

Many RNA species are modified post-transcriptionally to contain non-canonical bases, a process known as RNA editing. The most prevalent example of RNA editing is adenosine-to-inosine editing^1^, wherein adenosine deaminases (e.g., ADARs) enzymatically modify an adenosine base to an inosine base, which preferentially binds with cytosine bases rather than thymine (**Figure 1a**). The biological importance of adenosine-to-inosine editing is evident by the extreme phenotypes experienced as a result of perturbations to adenosine deaminases, such as defects in hematopoiesis^2^ and in neurological function^3^. It has been widely speculated that adenosine-to-inosine RNA editing may be a mechanism for nuclear retention and for other subcellular localization-based forms of post-transcriptional regulation of edited transcripts^4–7^. However, no previously existing methods have allowed for direct imaging of edited transcripts with single-molecule resolution, thus making it difficult to answer such questions. With that in mind, we developed inosineFISH (inoFISH), a fluorescence *in situ* hybridization-based method for directly imaging adenosine-to-inosine RNA editing events with single-molecule resolution.

**Figure 1.**
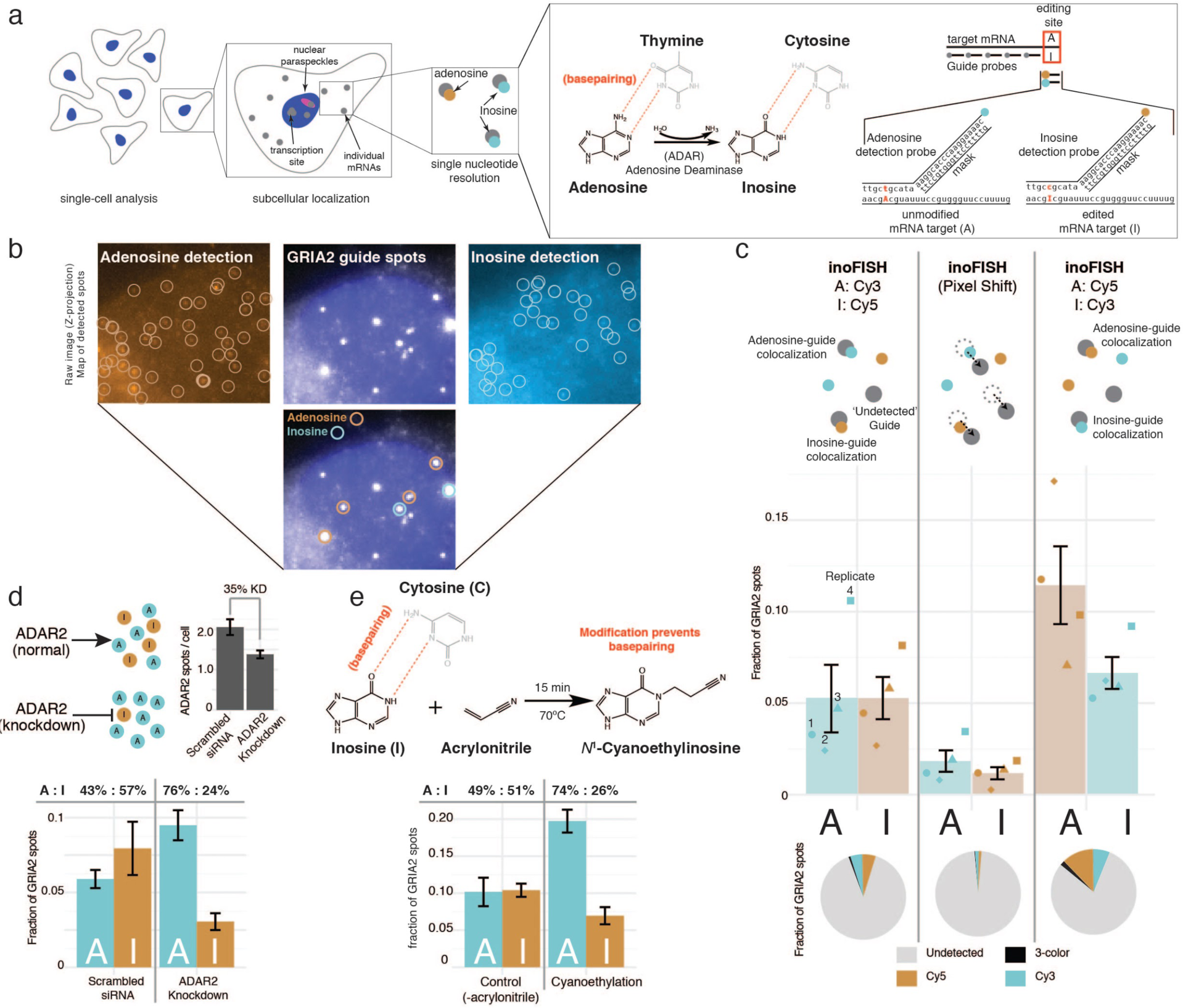
Fig 1: inoFISH specifically identifies individual adenosine-to-inosine edited transcripts *in situ*. (a) inoFISH probe hybridization scheme enabling discrimination between adenosine and inosine at editing sites in individual transcripts *in situ*. (b) Fluorescent spots called for *GRIA2* guide (middle), adenosine-detection (left) and inosine-detection (right) probes coupled to three different dyes colocalized (Overlay) to identify unedited and edited transcripts. (c) *GRIA2* inoFISH results from n = 4 SH-SY5Y biological replicates, including pixel-shift and dye-swap controls; schematic (top), inoFISH probe detection efficiencies per-replicate (points) and mean +/− s.e.m. (bars/error bars). Full summary of guide spot labels (pies); slice area as mean value over all replicates. (d) Schematic and *ADAR2* transcript abundance upon siRNA-mediated *ADAR2* knockdown (top) and *GRIA2* inoFISH results in SH-SY5Y cells (n = 3 replicates) after transfection with *ADAR2* siRNA and scrambled siRNA control (bottom). (e) Inosine cyanoethylation reaction schematic (top) and *GRIA2* inoFISH (bottom) results in SH-SY5Y cells (n = 3 inoFISH replicates) after *in situ* cyanoethylation with ‐acrylonitrile control.

Direct imaging of endogenous, edited RNA has been a challenge because single-base discrimination is difficult. Imaging techniques such as RNA fluorescence *in situ* hybridization (RNA FISH) rely on the hybridization of oligonucleotide probes to visualize the target of interest^8^. The challenge with using such a technique for visualizing edited RNAs is designing an oligonucleotide that is specific; short oligonucleotides will bind everywhere nonspecifically, but long oligonucleotides will not discriminate single-base differences. To circumvent these issues, we adapted a previously reported ‘toehold probe’ strategy^9^ to reduce the hybridized region of our detection probes in order to confer selectivity based on single-nucleotide differences in the target RNA (**Figure 1a**). In our scheme, one of the two competing detection probes targeted the unedited, adenosine-bearing sequence, and the other targets the edited, inosine-bearing sequence. We exploited the unusual base-pairing properties of inosine and treat these sequences as if they bear a guanosine (**Figure 1a**), thus enabling our probes to distinguish between adenosine and inosine. However, even when utilizing this ‘toehold probe’ strategy, single oligonucleotides are prone to nonspecific binding, so we simultaneously used a set of many oligonucleotide probes (the ‘mRNA guide’ probe) that target the constant, unedited portion of the mRNA, coupled to a different fluorophore than either of the SNV-detection probes (**Figure 1a**). The use of multiple oligonucleotides in the guide probe amplifies specific signal over non-specific binding of individual probes and showed us where to look for the detection probes.

We wanted to test whether inoFISH works for visualizing adenosine-to-inosine editing in a well-studied example, so we looked at Glutamate receptor 2 transcript (*GRIA2*) wherein the sequence that encodes for glutamine is edited to encode for arginine at amino acid position 607^10^. (*GRIA2* editing is critical for neuronal function^11^ and defects in *GRIA2* editing have been associated with ALS^12^.) RNA sequencing and restriction endonuclease digestion have shown *GRIA2* to be approximately 50% edited on average in SH-SY5Y cells^12^. We independently confirmed that *GRIA2* was edited by comparing the genomic DNA to the cDNA sequence in our SH-SY5Y cells (**Supplementary Fig. 1**). Then, we verified that the *GRIA2* guide spots observed by single molecule FISH were specific (**Supplementary Fig. 2**). Combining four biological replicates, 10.53% of the mRNA guides uniquely colocalized with adenosine or inosine detection probes, and we observed similar ratios of adenosine and inosine detection (a mean of 5.25% of *GRIA2* guides colocalized with the adenosine-detection probe and 5.28% of guides colocalized with the inosine-detection probe respectively; **Figure 1 b, c**), consistent with the previously reported ratios. To confirm that the detection probes are not colocalizing with the guide probes by random chance, we checked the frequencies at which randomly-placed guide spots would colocalize with the more abundant detection spots. In order to do this without violating the 3D shape of the cell, we computationally shifted spots in the guide channel images by 5 pixels in the X and Y planes (“Pixel shift”), thereby randomly repositioning the guide spots by moving them outside the range of any true colocalization events (see Methods; **Figure 1c, Supplementary Fig. 5**). From the pixel shift analysis, we measured 1.83% colocalization with adenosine and 1.16% colocalization with inosine, suggesting that most of the colocalization events we observed are in fact specific. To control for potential dye-specific effects such as differences in affinity for the target sequence, we swapped the fluorophores on the detection probes and repeated our analysis; **Figure 1c, Supplementary Fig. 5**).

To validate our measurements, we estimated the ratios of adenosine to inosine using three traditional, RT-PCR-based methods (**Supplementary Fig. 3**). First, we reverse transcribed *GRIA2* mRNA from SH-SY5Y cells and performed Sanger sequencing of the cDNA to estimate the ratio of adenosine to inosine using the relative height of the bases in the sequence chromatograph. We also performed a *BbvI* restriction digestion, which has a cut site that only appears in the cDNA when an adenosine-to-inosine event occurs and thus, provides an estimate of the fraction of transcripts that were edited based on the size of the digestion products as determined by bioanalyzer. Third, to determine the fraction of *GRIA2* cDNA coming from edited transcripts, we cloned and sequenced individual *GRIA2* cDNA molecules at the editing site. We observed the fraction of edited *GRIA2* transcripts in SH-SY5Y cells to be approximately 59%, 54.9% and 50% by Sanger sequencing, restriction digest and clonal analysis of cDNA respectively. These fractions of edited transcripts are consistent with the 49.86% fraction of edited transcripts observed by inoFISH. Further, we collected publicly available RNA-sequencing data from untreated SH-SY5Y cells^13^, and estimated fraction of edited transcripts by calculating the fraction of reads mapping to the editing site that call the editing site base as guanosine (see Methods and **Supplementary Fig. 3**).

While these results showed quantitative congruence between inoFISH and other methods, we needed to verify that inoFISH signals were specific to inosine bases and not adenosine or guanosine bases. We did this by altering the frequency of detected inosines in two different ways. First, we used an siRNA knockdown of ADAR2, the enzyme responsible for editing *GRIA2* transcripts, in order to lower the fraction of *GRIA2* editing transcripts in SH-SY5Y cells^14,15^. We observed 57.41% estimated editing in the scrambled siRNA control, compared with 24.33% estimated editing in the ADAR2 knockdown samples (the total number of *GRIA2* transcripts remained equal) (**Figure 1d**). This result shows that inoFISH can discriminate between adenosine and inosine bases. Next, we adapted a biochemical tagging method, cyanoethylation, whereby inosine bases are modified with acrylonitrile by Michael addition^16,17^. This modification adds acrylonitrile to the N^1^ position of the inosine base and prevents base pairing to cytosine. Using this method, we significantly reduced the percentage of transcripts that we called as edited from 50.5% to 26.1% (**Figure 1e, Supplementary Fig. 6**). Note that a mild cyanoethylation treatment was used in order to make the method compatible with inoFISH, rendering the inosine conversion incomplete. This result demonstrated that inoFISH specifically detects inosine-bearing transcripts rather than guanosine.

We next measured subcellular localization of edited and unedited transcripts. One example of subcellular localization is nuclear retention of edited mRNAs. Previous studies used cell fractionation, RT-PCR and thin-layer chromatography to show that unmodified RNAs existed in both cellular compartments, whereas hyper-edited RNAs—but not selectively edited RNAs—were retained in the nuclear fraction^4,6^. Studies from other groups have also suggested that adenosine-to-inosine editing is important for RNA localization by showing that mRNAs containing Alu repeats, which are prone to adenosine-to-inosine editing, are inefficiently exported to the cytoplasm^5^.

With this in mind, we looked for any association between editing status and subcellular localization of *GRIA2* transcripts, which are selectively edited. We determined whether guide spots were in the nucleus by seeing if they overlapped with the nuclear stain DAPI. We observed roughly equal fractions of edited to unedited *GRIA2* transcripts in both cellular compartments (cytoplasm: 52.9%, nucleus: 48.3%); **Figure 2a**). (Surprisingly and uncharacteristically of mRNAs in general, we also found that 93.4%% of *GRIA2* transcripts localize to the nucleus in SH-SY5Y cells (**Supplementary Fig. 2**), irrespective of editing. Despite this strong localization, the *GRIA2* transcript was still translated and we explore this result further in **Supplementary Fig. 2**.)

**Figure 2.**
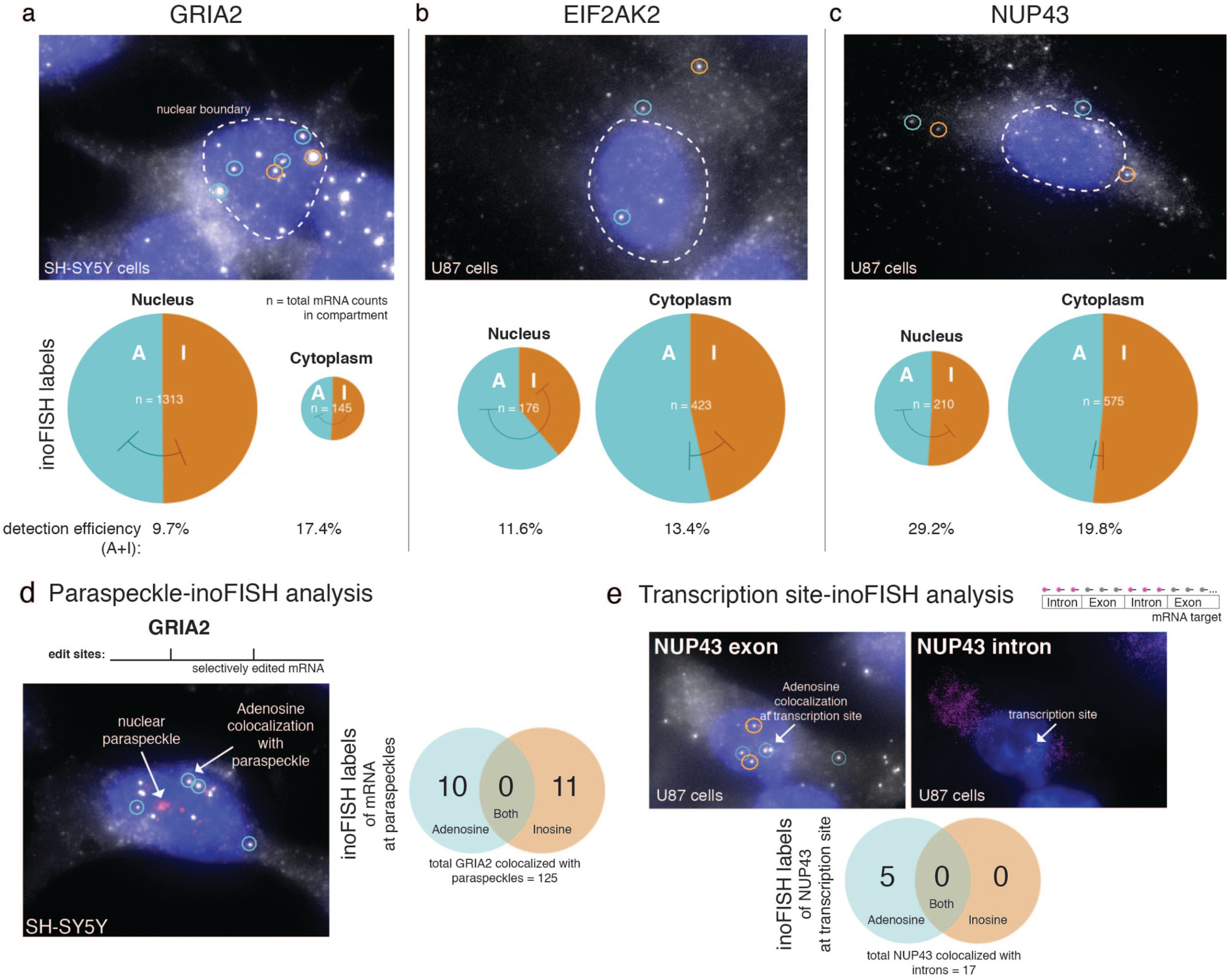
Fig. 2: inoFISH reveals that editing is not important for subcellular localization of *GRIA2* in SH-SY5Y cells, or for *EIF2AK2*, and *NUP43* in U-87 MG cells. inoFISH results with nuclear localization analysis of (a) *GRIA2* (n = 4 replicates), (b) *EIF2AK2* (n = 3) and (c) *NUP43* (n = 2) transcripts: representative overlays (top) and fractions of labeled transcripts found to be unedited or edited (bottom). (d) inoFISH results with NEAT1-colocalization for paraspeckle localization analysis of (left) GRIA2 (n = 4) and (right) *EIF2AK2* (n = 3): schematic and representative overlays (top) and counts of inoFISH-labeled, paraspeckle-associated transcripts (bottom). (e) *NUP43* inoFISH results with transcription site localization analysis (n = 2): schematic and representative images (top) and counts of inoFISH-labeled, transcription site-associated *NUP43* transcripts (bottom).

We also applied inoFISH to visualize adenosine-to-inosine editing in two additional targets: the hyperedited transcript *EIF2AK2*^18^ (**Figure 2b**) as well as the Alu-bearing *NUP43*^5^ (**Figure 2c**) in U87-MG cells (**Supplementary Fig. 4**) that we independently validated for editing by comparing the genomic DNA sequence to the cDNA sequence (**Supplementary Fig. 1**). Using inoFISH, we found that 6.91% of *EIF2AK2* guide spots colocalized with the adenosine-specific detection spots and 5.57% of guide probe colocalized with inosine-specific detection spots (**Figure 2b, Supplementary Fig. 5**), giving a population-wide editing rate estimate of 44.6%. For *NUP43*, 11.3% of guide spots colocalized with the adenosine-specific detection spots and 12.4% of guide probe colocalized with inosine-specific detection spots (**Figure 2c, Supplementary Fig. 5**), giving a population-wide editing rate estimate of 47.7%. As with GRIA2, these results are consistent with traditional measurement methods for determining fraction of edited transcripts, such as RT-PCR with Sanger sequencing for *EIF2AK2* and *NUP43* in U-87 cells **(Supplementary Fig. 1)**^19^. We biochemically tested for specificity of our inosine-detection probe by cyanoethylation and found that cyanoethylation reduced the percentage of inosine-detection probe colocalization with guide probe for both *EIF2AK2* and *NUP43*, showing that the method is specific (**Supplementary Fig. 6**).

InoFISH also allowed us to test the suggestion from previous studies that edited transcripts may be trafficked to nuclear paraspeckles^7^. To test whether selectively-edited *GRIA2* localized to paraspeckles, we performed inoFISH together with single-molecule RNA FISH of *NEAT1* RNA, a marker of nuclear paraspeckles^20^, in SH-SY5Y cells and U87-MG cells respectively (**Figure 2d**). We first checked whether there was association of any *GRIA2* transcripts (irrespective of editing status) with paraspeckles; we found that 8.57% of all *GRIA2* transcripts colocalize with paraspeckles in SH-SY5Y cells (**Figure 2d**). To test whether transcript-paraspeckle association was greater than one would expect by chance, we computationally simulated the distribution in the case of purely random paraspeckle associations for each experiment, given each observed cell’s (1) shape of nucleus, (2) number of transcripts retained in the nucleus, and (3) location of all paraspeckles (**Supplementary methods**). We found that the observed rate of *GRIA2*-paraspeckle association is in fact 1.70 fold greater than one would expect by random chance (representative p < 0.001).

We then used inoFISH to further interrogate whether edited or unedited *GRIA2* transcripts were preferentially found in the observed transcript-paraspeckle associations. We computationally simulated the distributions in the case of purely random association rates of paraspeckles with edited and unedited transcripts (**Supplementary methods**). We found no significant differences in the editing status in paraspeckles for *GRIA2* in SH-SY5Y cells (representative p = 0.44; **Figure 2d**). This result demonstrates that paraspeckle association of edited transcripts is not universal.

Next, we wanted to apply our new tool to determine whether adenosine-to-inosine editing is co-transcriptional or post-transcriptional. We screened *NUP43* in U-87 MG cells for editing status at the transcription site by concurrently performing *NUP43* inoFISH with single-molecule FISH targeting *NUP43* introns (**Figure 2e**). Introns are markers of transcription sites by single-molecule FISH, and colocalization of an edited transcript with intron signal would suggest that editing can be a co-transcriptional process. After screening 212 cells, in which we observed 17 total transcription sites, 5 transcription sites were labelled as containing unedited *NUP43* and none were labelled as containing edited *NUP43* (**Figure 2e**). This result does not rule out co-transcriptional editing of *NUP43* altogether, but it does at least suggest that *NUP43* editing may be post-transcriptional.

We also looked for evidence of fluctuations in editing rate from cell to cell. We modeled what it would look like if single cells exhibited constant editing rates similar to the population average (e.g. a constant 50% single-cell editing rate in all cells; **Figure 3a**), and what it would look like if single cells had fluctuations in editing rates (e.g., in the extreme case that some cells have essentially only unedited or only edited transcripts; **Figure 3b**). Next, we calculated the observed counts of edited and unedited transcripts for *GRIA2* and *NUP43* on the single-cell level and found that *GRIA2* editing is not consistent with the constant-rate model, suggesting that the single-cell rate of editing is not constant (**Figure 3c**). We also found that *NUP43* editing was consistent with the constant-rate model in U-87 MG cells (**Figure 3d**). These results suggest that single-cell fluctuations in the rate of editing may occur in a target-dependent, cell-type specific manner.

**Figure 3.**
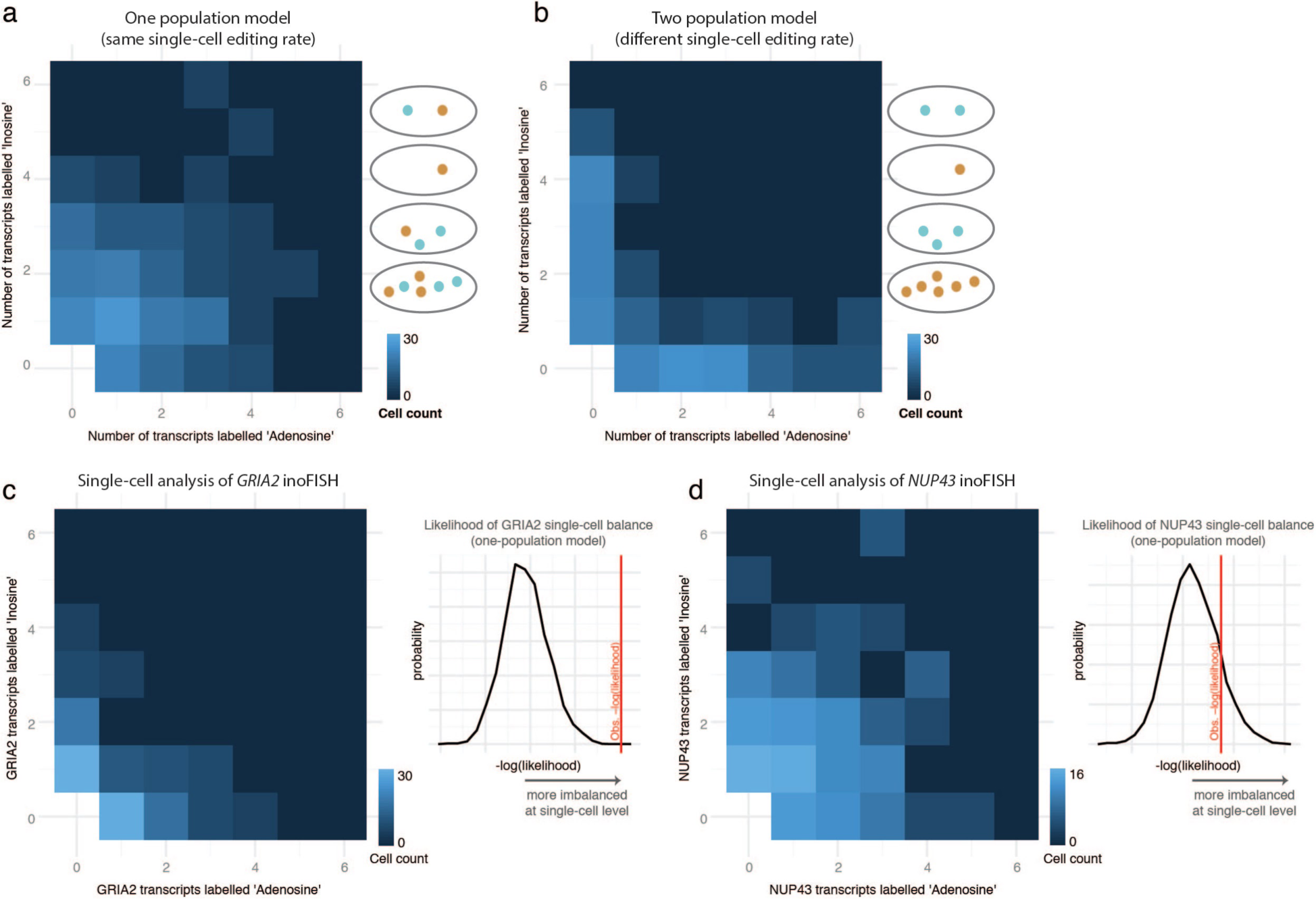
Fig. 3: Single-cell analysis of inoFISH results uncovers transcript-specific variability in single-cell editing rates. (a) Simulated inoFISH results assuming binomially-distributed per-cell counts of edited and unedited transcripts ~ *Bin(n, p)*, conditioned on *p,* the observed population-wide *NUP43* editing rate and *n* the number of labelled transcripts for each cell in the *NUP43* dataset. (b) Simulated inoFISH results assuming two populations of cells; one population with 95% editing and the other with 5% editing mixed in proportion according to *p*. (c) Single-cell analysis of *GRIA2* inoFISH results pooled over all 4 replicates (left) and simulation of the exact conditional null distribution of −log(likelihood) of the data under the binomial model specified in (a) (right), but conditioned on *p,* the observed population-wide *GRIA2* editing rate and *n* the number of labelled transcripts for each cell in the *GRIA2* dataset (d) Single-cell analysis of *NUP43* inoFISH results pooled over all 4 replicates (left) and simulation of the exact conditional null distribution of −log(likelihood) of the data under the binomial model specified in (a) (right).

InoFISH provides a direct method to visualize adenosine-to-inosine RNA editing in single cells with single-nucleotide resolution. Cell population-wide studies lack the resolution to provide information such as subcellular localization and cell-to-cell variability of RNA editing. This new tool will enable researchers to answer basic questions about edited RNA species and will enable a deeper understanding of the biology of adenosine-to-inosine editing.

## Materials and Methods

### Cell culture

We grew human neuroblastoma cells (SH-SY5Y, ATCC CRL-2266) in a 1:1 mixture of Eagle’s Minimum Essential Medium and F12 Medium supplemented with 10% FBS and 50 U/mL penicillin and streptomycin. We grew human glioblastoma cells (U-87 MG, ATCC HTB-14) in Eagle’s Minimum Essential Medium supplemented with 10% FBS and 50 U/mL penicillin and streptomycin. Note: Some SH-SY5Y and U87 cells can be autofluorescent when grown on glass slides, this is improved when the cells are ~70% confluent.

### Selection of targets for inoFISH

Besides the well-studied editing target *GRIA2*, we wanted to test inoFISH on editing targets that are less commonly studied in the literature. For these we referred to the literature, to the RADAR database of RNA editing, and to publicly available RNA-seq data for screening. We identified *NUP43* and *EIF2AK2* as targets that are studied in a relatively small number of research groups, that are candidate editing targets in published transcriptome-wide adenosine-to-inosine editing screens, and that have editing sites amenable to inoFISH probe designs. In order to systematically identify inoFISH targets, we found it useful to first filter targets by those with enough non-repetitive sequence to design a full guide probe (>24 20-mer oligonucleotides). Then, we performed transcriptome-wide screens aimed at studying editing targets in the RADAR database that are conserved across humans, non-human primates, and mice^21^. We expect that conserved targets are more likely both to be biologically interesting and to be observable in our cell lines than targets that only appear in screening experiments in humans alone.

#### RNA-sequencing-based screen for candidate editing targets

We downloaded publicly available RNA-sequencing data from EBI ArrayExpress for total RNA from two biological replicates of SH-SY5Y cells^13^ and three replicates of U-87 MG cells^19^. We also downloaded RNA editing candidates’ positions (in human genome build hg19) from the RADAR database, with a focus on conserved positions and those derived from neural lineage samples. We aligned raw reads to hg19 with STAR v2.3.0e with default parameters except for runThreadN set to 4. Next, we used picard tools v1.96^22^ to interconvert between aligned SAM and BAM file formats (SamFormatConverter), to sort BAMs by coordinate (SortSam), to index sorted BAMs (BuildBamIndex), and to remove PCR duplicates (MarkDuplicates). We then trimmed 5’ ends of all aligned and PCR-deduplicated reads with bamUtil v1.0.13^23^. Finally, using bam-readcount v^24^ we filtered aligned reads by those overlapping candidate editing sites of interest, by their overall mapping (phred) score and individual base call qualities using a custom bed file and parameters −q 25 and −b 20. We quantified RNA-seq based editing rate estimates in R by calculating the fraction of filtered reads with ‘G’ base calls at each editing site. We considered for downstream RNA editing verification and inoFISH probe design any sites in these screens with at least seven overlapping filtered reads, of which at least one was called ‘G’ at the editing site.

#### Verification of RNA editing of candidate targets

As described below, we used RT-PCR of total RNA and PCR of genomic DNA in cell lines of interest to further check that candidate editing sites were in fact RNA edited in our cell lines.

#### Optimization of inoFISH targets

We then chose targets for inoFISH by checking RT-PCR and genomic DNA PCR Sanger sequencing results for each candidate site to ensure that there would be no additional polymorphisms in transcripts, resulting either from RNA editing or SNPs, in the regions flanking the editing site up to 30-bp up-or downstream. (**Supplementary Fig. 4**). We ultimately designed inoFISH probe sets against one editing site in *NUP43* and two editing sites in *EIF2AK2* (both using the same guide probe set) that appeared to be amenable to inoFISH guide and detection probe set design. We were able to verify inoFISH probe binding with detection efficiencies greater than expected by random colocalization for the one *NUP43* candidate editing site and one of the two EIF2AK2 editing sites. (See below for experimental methods.)

### Genotyping of edited regions

We extracted genomic DNA from SH-SY5Y cells and U-87 MG cells using the Qiagen DNeasy Blood & Tissue kit. We used Platinum Taq (Invitrogen cat. #10966-018) for PCR amplification of the genomic regions of interest for each target, following manufacturer’s recommendations for reaction component concentrations. We PCR-amplified two biological replicates, each with two technical PCR replicates. For *GRIA2*, we used primers *GRIA2*-F1 and *GRIA2*-R2 (**Supplementary table 1**). For *EIF2AK2*, we used *EIF2AK2*_20-F1 and *EIF2AK2*_20-R1, and for *NUP43* we used *NUP43*-F1 and *NUP43*-R1 (**Supplementary table 1**). We confirmed PCR product sizes by gel electrophoresis, using a 1.5% agarose gel in TAE. Then, we treated these PCR products with ExoSAP-IT (Affymetrix 78200) according to manufacturer’s instructions, and submitted them for Sanger sequencing at the University of Pennsylvania DNA Sequencing facility.

### Estimation of editing efficiency by RT-PCR and Sanger sequencing

We extracted total RNA from SH-SY5Y and U-87 MG cells using miRNeasy kits (Qiagen 217004) according to manufacturer’s instructions. Then, we reverse-transcribed target transcripts around editing sites of interest using Superscript III First strand RT kit (ThermoFisher 18080044) according to manufacturer’s instructions. In separate reactions for RNA from each cell type, we used both oligo-dT and transcript-specific primers for reverse-transcription. Briefly, we used 50 ng of RNA per reaction for reverse transcribed with either oligo-dT or transcript-specific primers (**Supplementary table 1**). Then, we performed PCR with transcript-specific primers (**Supplementary table 1**) using Platinum Taq (Invitrogen cat. #10966-018) according to manufacturer’s instructions. We completed biological replicates, each with technical PCR replicates for these reactions. We confirmed product sizes by gel electrophoresis on 1.5% agarose gels in TAE. Then, we treated these products with ExoSAP-IT (Affymetrix 78200) according to manufacturer’s instructions and submitted for sequencing at the University of Pennsylvania DNA Sequencing facility.

### Estimation of editing efficiency by clonal analysis of *GRIA2* RT-PCR product

The amplified *GRIA2* cDNA was cloned into a vector using the TOPO TA cloning kit (Thermo), transformed into chemically competent *Escherichia coli* cells, and plated on LB plates with 0.1 mg/mL ampicillin. We isolated DNA from >20 Individual colonies and submitted it for sequencing at the University of Pennsylvania DNA Sequencing Facility. We performed sequence alignment at the editing site using MAFFT in Benchling to determine the ratio of edited and unedited transcripts.

### Estimation of editing efficiency by restriction digest and bioanalyzer analysis

Edited and unedited *GRIA2* cDNAs yield distinct restriction fragment patterns upon digestion with *Bbv*I^25^. Edited *GRIA2* cDNA yields two DNA fragments upon digestion (225 bp and 46 bp), and unedited *GRIA2* cDNA yields three DNA fragments (145 bp, 80 bp, and 46 bp). Following *BbvI* digestion (NEB R0173S) of *GRIA2* cDNA, according to manufacturer’s instructions, we submitted digestion products for fragment sizing analysis on an Agilent 2100 Bioanalyzer at the University of Pennsylvania DNA Sequencing Facility.

### RNA probe design and synthesis

For each of the validated editing sites, we designed probes by matching free energies of hybridization as specified in Levesque et al. (2013b). We optimized mask oligonucleotides to leave 8-base-pair (bp) overhangs for each of the SNP probes and pooled all five together to act as the complete allele-specific probe. We provide all oligonucleotide sequences in Supplementary Table 1. We coupled 3’ amine-labeled adenosine-and inosine-detection probes to NHS-Cy3 or NHS-Cy5 fluorophores (GE Healthcare) and purchased respective guide probes labeled with Cal fluor 610 (Biosearch Technologies). We coupled probes targeting ADAR1, ADAR2 and NEAT1 mRNA to NHS-Atto700. We purified dye-coupled probes by high-performance liquid chromatography.

### inoFISH procedure

We grew cells on glass coverslides until ~70% confluent. We washed the cells twice with 1X PBS, then fixed for 10 minutes with 4% formaldehyde/1X PBS at room temperature. We aspirated off the formaldehyde, and rinsed twice with 1X PBS prior to adding 70% ethanol for storage at 4°C or inoFISH after a one hour permeabilization in 70% ethanol. We incubated our cells overnight at 37°C in hybridization buffer (10% dextran sulfate, 2× SSC, 10% formamide) with 100 nM concentration of guide probe, 24 nM concentration of the adenosine‐ and inosine-detection probes and 72 nM concentration of the mask probe, ensuring excess mask for complete hybridization to the detection probes. The following morning, we performed two washes in wash buffer (2X SSC, 10% formamide), each consisting of a 30-min incubation at 37°C. After the second wash, we rinsed once with 2X SCC/DAPI and once with anti-fade buffer (10 mM Tris (pH 8.0), 2X SSC, 1% w/v glucose). Finally, we mounted the sample for imaging in an anti-fade buffer with catalase and glucose oxidase (Raj et al. 2008) to prevent photobleaching. We performed RNA FISH on cell culture samples grown on a Lab-Tek chambered coverglass using 50 μL of hybridization solution spread into a thin layer with a coverslip and placed in a parafilm-covered culture dish with a moistened Kimwipe to prevent excessive evaporation.

### Imaging

We imaged each samples on a Nikon Ti-E inverted fluorescence microscope using a 100× Plan-Apo objective (numerical aperture of 1.40) and a cooled CCD camera (Andor iKon 934). For 100× imaging, we acquired z-stacks (0.3 μm spacing between stacks) of stained cells in five different fluorescence channels using filter sets for DAPI, Cy3, Calfluor 610, Cy5, and Atto 700. The filter sets we used were 31000v2 (Chroma), 41028 (Chroma), SP102v1 (Chroma),17 SP104v2 (Chroma) and SP105 (Chroma) for DAPI, Atto 488, Cy3, Atto 647N/Cy5 and Atto 700, respectively. A custom filter set was used for Alexa 594/CalFluor610 (Omega). We tuned the exposure times depending on the dyes used: 4 seconds for each guide probe, 4000 msec for each of the detection probes, 5000 msec for the NEAT1 probe, and 7000 msec for ADAR1 and ADAR2 probes. We also acquired images in the Atto 488 channel with a 1000 msec exposure as a marker of autofluorescence.

### Image analysis

We first segmented and thresholded images using a custom Matlab software suite (downloadable at https://bitbucket.org/arjunrajlaboratory/rajlabimagetools/wiki/Home). Segmentation of cells was done manually by drawing a boundary around non-overlapping cells. The software then fits each spot to a two-dimensional Gaussian profile specifically on the Z-plane on which it occurs in order to ascertain subpixel-resolution spot locations. Colocalization took place in two stages: In the first stage, guide spots searched for the nearest-neighbor SNP probes within a 2.5-pixel (360-nm) window. We ascertained the median displacement vector field for each match and subsequently used it to correct for chromatic aberrations. After this correction, we used a more stringent 1.5-pixel (195-nm) radius to make the final determination of colocalization. In order to test random colocalization due to spots occurring randomly by chance, we took our images and shifted the guide channel by adding 5 pixels (1.3 μm) to the X and Y coordinates and then performing colocalization analysis.

### Autofluorescence subtraction

For U-87 MG cells, we controlled for punctate autofluorescence by imaging with the 41028 (Chroma) filter set, the ‘gfp channel’, which we have previously found to be sensitive for autofluorescence in this cell line (data not shown). We performed colocalization as previously described between guide spots and any spot-like autofluorescence called in the gfp channel. In R, we excluded spots colocalizing with this autofluorescence from all inoFISH analyses.

### Subcellular localization

#### Nuclear localization

We extracted a DAPI nuclear mask as previously described^8^. We call a spot as localized to the nucleus if the guide spot X and Y coordinates overlap with the 2D nuclear mask. *Localization to transcription sites*. We visualized *NUP43* introns by probing with intron-specific probes coupled to Atto 700 and imaging with SP105 filter set. We used the txnSiteGUI2 interface within rajlabimagetools to manually curate calls of exon-intron spot colocalization. *Localization to paraspeckles.* We visualized paraspeckles by probing with NEAT1-specific probes coupled to Atto 700 and imaging with SP105 filter set. We used the txnSiteGUI2 interface within rajlabimagetools to manually curate calls of transcript-paraspeckle association.

### In situ cyanoethylation

Cyanoethylation was performed similarly to previous descriptions^16,17^. We aspirated the 70% ethanol off of the fixed cells and added cyanoethylation solution (1.1 M triethylammoniumacetate (pH 8.6) resuspended in 100% ethanol) with or without 1.6 M acrylonitrile at 70 °C for 15 min. Use large volume to prevent drying from evaporation. Remove from heat after 15 min (30 min incubation abolishes guide probe signal) and wash twice with wash buffer (2× SSC, 10% formamide) before beginning inoFISH procedure.

### Statistical analysis

#### Detection efficiency

For each label (edited or unedited) in each experiment we calculated the mean fraction of transcripts colocalized with a spot of that label over all replicates (excluding 3-color spots). We also calculated the standard error of this mean, plotted on each bar chart as the magnitude of error bars in one direction and on each pie plot as the scaled radial error bar internal to each slice.

#### Population-wide editing rate estimation by inoFISH

We define the population-wide editing rate estimate as the average over all replicates of the fraction of uniquely labelled guide spots labelled as edited. Formally, for *e,* the mean fraction of guide spots colocalized with an editing-detection spot, and *u,* the mean fraction of guide spots colocalized with an unedited-detection spot, the population-wide estimated editing rate *p_e_* is *p_e_* = *e*/(*e* + *u*).

#### Paraspeckle-transcript association rates

In MATLAB, we simulated the exact conditional null distribution of paraspeckle-transcript association rates for each experiment under the null hypothesis that a paraspeckle and a nuclear-localized transcript will only colocalize by chance. For each cell in each experiment, we conditioned on (1) the shape and size of that cell’s nucleus, (2) the locations of all paraspeckles in that nucleus, and (3) the number of transcripts of interest (*GRIA2* or *EIF2AK2*) retained in the nucleus. In order to efficiently simulate these distributions, rather than using txnSiteGUI2 as above, we generated 2D masks for paraspeckle locations and called paraspeckle-transcript association when a randomly placed transcript spot overlapped with this mask. We selected the mask size as 25 pixels per paraspeckle spot called—roughly the mean paraspeckle size—based on our inspection of paraspeckles while calling spots (as in Image analysis). For each experiment, we simulated draws from the exact conditional null distribution 1000 times. A raw p-value for paraspeckle-transcript association rate is equal to the fraction of simulations with a higher paraspeckle-transcript association rate. We similarly simulated exact conditional null distributions for paraspeckle-edited-transcript and paraspeckle-unedited-transcript association rates.

#### Single-cell editing rate distributions

In R, we assessed single-cell spot counts after inoFISH colocalization as reported by rajlabimagetools (in Image analysis), as well as after autofluorescence subtraction (for U-87 MG data). We simulated the null distribution of data likelihood under a null model wherein all cells sharing the same effective editing rate: for an experiment with overall estimated editing rate equal to *p_e_* (above), let n_e_^j^ be the number of edited transcripts detected in cell j and n_u_^j^ be the number the number of unedited transcripts detected in cell j. Under the null model, n_e_^j^ is drawn from a Binomial with (n_e_^j^ + n_u_^j^) draws and probability *p_e_*. We simulated single-cell label counts for cells by drawing from these conditional null distributions for each cell 100000 times. We then compared the negative log-likelihood of the observed data, combined over all replicates, with the distribution of negative log-likelihoods of each simulation iteration. A p-value of 0.12 indicates that 12% of the simulated iterations had a negative log-likelihood that was greater than the observed data. Note that the −log(likelihood) density plots in Fig. 3 are subsampled to 3000 of the aforementioned 100000 such iterations per plot, in order to facilitate figure generation.

### siRNA knockdowns of ADAR2

Briefly, we used Lipofectamine RNAimax to transfect SH-SY5Y cells with Silencer Select siRNAs targeting ADARB1 (ADAR; ID:s1011, Ambion) and a Negative Control siRNA (#1, Ambion) for 72 hrs, verifying knockdown via RNA FISH of ADAR2. **Cell cycle inhibitor.** We measured nuclear retention of *GRIA2* mRNA by inhibiting transcription for 24 hr by applying aphidicolin at 1 ug/ml.

### Reproducible analyses

Scripts for all analyses presented in this paper, including all data extraction, processing, and graphing steps are freely accessible at the following url: https://www.dropbox.com/sh/qj2hanxe3hs14ry/AACx_3IO9LVVYX4DhKWxqoKla?dl=0.

## Acknowledgements

SHR, IAM and AR conceived of the paper, SHR and IAM performed experiments, IAM, RG, and AR wrote custom software, IAM, SHR and AR analyzed the data, IAM, SHR and AR wrote the paper. We thank Drs. Nishikura and Sakurai for helpful discussions, as well as Caroline Bartman and Benjamin Emert for comments and suggestions. IAM acknowledges support from NIH/NIGMS T32GM007170 (University of Pennsylvania MSTP), NIH/NHGRI T32HG000046 and NIH/NINDS F30NS100595 and AR from an NSF CAREER Award 1350601, NIH New Innovator 1DP2OD008514, NIH/NIBIB R33 EB019767, and NIH 4DN U01 HL129998.

